# Simulating pedigrees ascertained on the basis of observed IBD sharing

**DOI:** 10.1101/2024.05.13.594012

**Authors:** Ethan M. Jewett, the 23andMe Research Team

## Abstract

In large genotyping datasets, individuals often have thousands of distant cousins with whom they share detectable segments of DNA identically by descent (IBD). The ability to simulate these distant relationships is important for developing and testing methods, carrying out power analyses, and performing population genetic analyses. Because distant relatives are unlikely to share detectable IBD segments by chance, many simulation replicates are needed to sample IBD between any given pair of distant relatives. Exponentially more samples are needed to simulate observable segments of IBD simultaneously among multiple pairs of distant relatives in a single pedigree. Using existing pedigree simulation methods that do not condition on the event that IBD is observed among certain pairs of relatives, the chances of sampling shared IBD patterns that reflect those observed in real data ascertained from large genotyping datasets are vanishingly small, even for pedigrees of modest size. Here, we show how to sample recombination breakpoints on a fixed pedigree while conditioning on the event that specified pairs of individuals share at least one observed segment of IBD. The resulting simulator makes it possible to sample genotypes and IBD segments on pedigrees that reflect those ascertained from biobank scale data.

## 2. Introduction

Simulations of genetic transmission within fixed pedigrees are used for a many applications, such as generating training or testing sets to support method development ^8,11,20,22,23^, performing power analyses ^4,5,16,21^, studying biological phenomena or processes such as inbreeding ^12,17^, and empirically obtaining probability distributions that are used for relationship inference ^2,9^.

Because the probability that two distant relatives share tracts of DNA identically by descent (IBD) can be small, the probability of sampling observed tracts of IBD when simulating a pair of distant relatives can also be small. For example, fourth cousins have only a fifty percent chance of sharing a detectable segment of IBD^9,26^ while sixth cousins share detectable IBD segments less than five percent of the time ^9,26^, even when using realistic simulations that account for cross-over interference and sex-specific genetic maps ^3^. Moreover, the probability of sharing IBD decreases exponentially quickly as the degree of the relationship increases ^6,9,26^.

When multiple distant relationships are considered simultaneously within a pedigree, the probability of sampling detectable IBD among all relative pairs can be very low. For example, given that fourth cousins share detectable IBD less than half the time, the probability of sampling IBD in a pedigree with just ten fourth degree relative pairs is less than 1/2^10^ ≈ 10^*−*4^ if the pairs are approximately mutually independent. For ten sixth cousins, the probability is less than 1/20^10^ ≈ 10^*−*13^.

Although the probability of IBD sharing through a particular pair of common ancestors is small, individuals have many distant relatives with whom they share many common ancestors. Thus, individuals in biobank-scale datasets typically have thousands of fourth and fifth cousins ^9^ with whom they share detectable amounts of IBD. The ability to simulate genetic transmission on pedigrees relating subsets of such distant relatives is important for testing methods that aim to infer these pedigrees ^11,20,23^ or which use the IBD among distant relatives for downstream analyses ^8,19^.

In pedigrees in which relatives are closely-related, one can use an existing simulation method together with rejection sampling to obtain samples under the conditional distribution by rejecting any sample in which the required individuals share no IBD. However, when relatives are distantly related, rejection sampling becomes impractical because the fraction of rejected samples is close to one. Therefore, it is necessary to simulate using a method that allows pedigree sampling conditional on the event that particular pairs share IBD. Although many methods have been developed for simulating meioses within a pedigree structure ^1,3–5,13,15,16,18,21,24,25^, no existing method makes it possible to condition on the event that two or more individuals share IBD.

Here, we show how to sample meioses and transmitted IBD among pairs of individuals and within a pedigree, conditional on the event that two or more individuals share IBD with one another. We then demonstrate the utility of this simulator for two downstream applications: pedigree inference and the simulation of an individual together with their set of distant relatives.

## 3. Sampling conditional on any observed IBD

Suppose two individuals (*i* and *j*) are related through relationship *R* and suppose we know that they share some nonzero, but unspecified, amount of IBD I. We may simply know that *I* is greater than zero, or we may know that *i* and *j* share at least one segment longer than a minimum observable length threshold τ . We want to simulate all recombination events on the ancestral path connecting *i* and *j*, conditional on the event that *I* > 0 or conditional on the event that they share at least one segment longer than τ . We then want to scale this up to sample meioses in a pedigree in which arbitrary sets of individuals are known to share IBD with one another.

In this paper, we will use the notation of Ko and Nielsen ^14^ to specify a relationship *R* = (*u, d, a*) between *i* and *j*, where *u* is the number of meioses “up” from *i* to their common ancestor(s) with *j* and *d* is the number of meioses “down” from the common ancestor(s) to *j*. The number of common ancestors is *a*, which is either 1 or 2 in an outbred pedigree. Although inbreeding is common in many populations and although all human pedigrees contain loops when we consider a sufficient number of generations in the past, we will focus on outbred pedigrees here because it makes the bookkeeping simpler. The model can be extended to include consanguineous relationships by including considerably more bookkeeping, but the method is also slower in the case of inbreeding because of the need to sample larger groups of meioses at the same time to satisfy constraints.

We want to sample all *u* meioses up from *i* to its first common ancestor with *j* and all *d* meioses down from its first common ancestor to *j* (black meioses in Figure 1). If *i* and *j* have two common ancestors (a = 2) then we must sample two additional meioses (orange meioses in Figure 1) leading to their second shared common ancestor.

**Figure 1.**
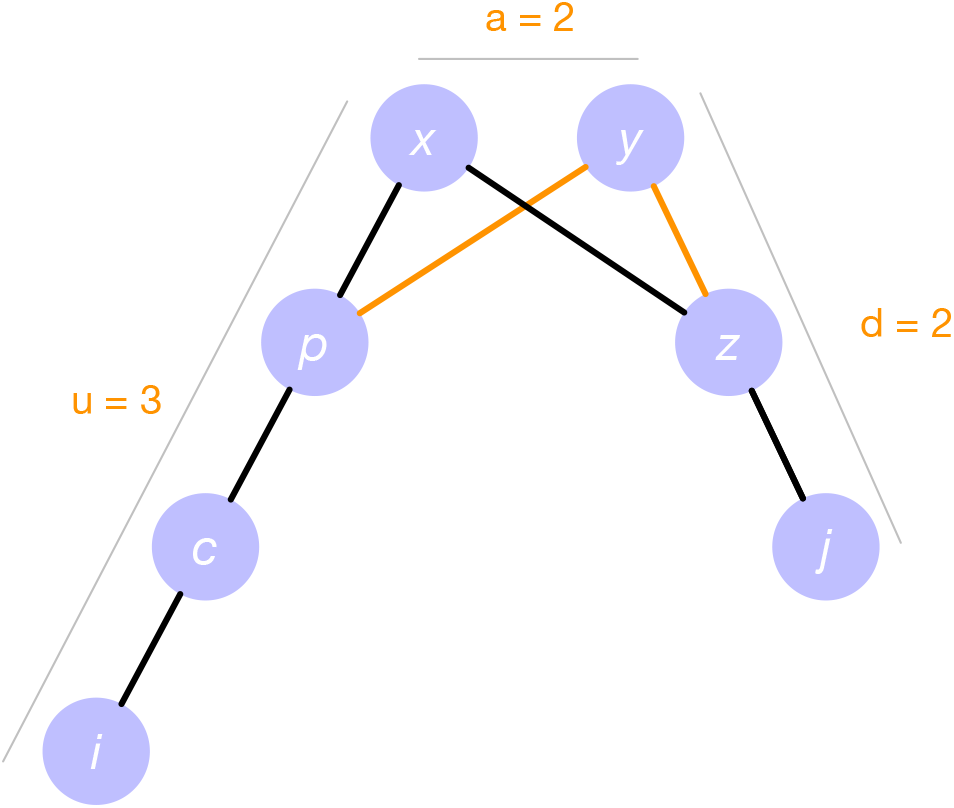
A pedigree of individuals (circles) with meioses between them (lines). Nodes *i* and *j* are related through common ancestors *x* and *y*. The number of meioses “up” from *i* to its common ancestors with *j* is *u* = 3. The number of meioses down from the common ancestors to *j* is *d* = 2. The number of common ancestors is *a* = |{*x, y*}| = 2. This yields the relationship *R* = (3, 2, 2) between *i* and *j* using the notation of Ko and Nielsen ^14^.

### 3.1. IBD propagation

In a pedigree, genetic material is passed only from ancestors to descendants. However, in developing the sampling framework presented in this paper, it is useful to conceptualize the transmission of tracts of IBD rather than genetic material itself. A tract of IBD is a contiguous region along a single chromosome that is shared identically by descent between two individuals. For example, in Figure 2, the blue regions depict the tracts of IBD shared between an individual and each of their ancestors of various degrees.

**Figure 2.**
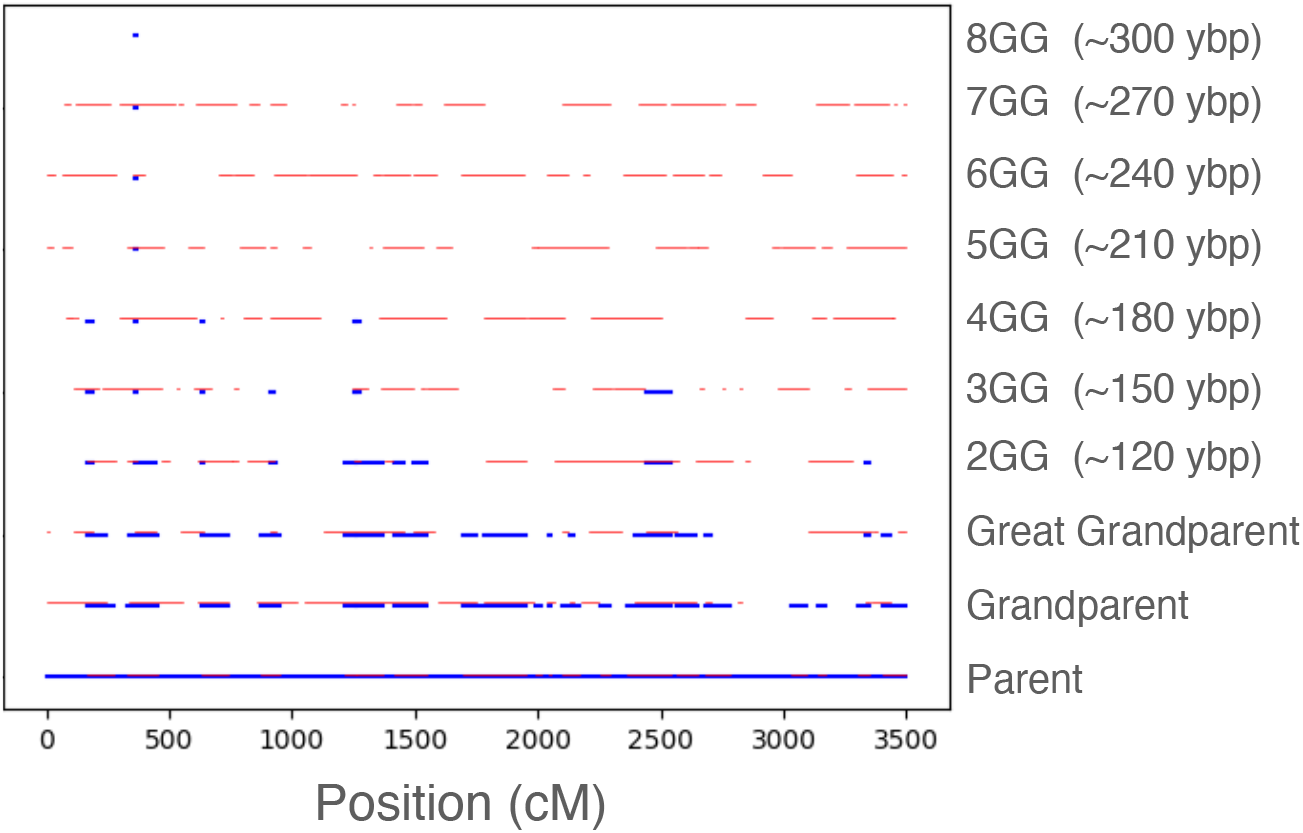
One unconditional simulation replicate showing the tracts of IBD transmitted between relatives of various degrees. Each row shows the tracts of IBD (blue) that are shared between an individual (not shown) and an ancestor along the full linear genome. The bottom row shows that the individual shares exactly one copy of their full linear genome (half their entire diploid genome) with a parent. The second line from the bottom shows that they share approximately half their linear genome (1/4 of their diploid genome) with a grandparent. Thin red lines show the regions between recombination breakpoints that are shared between adjacent ancestors (e.g., between a 3^*rd*^ great grandparent and a 4^*th*^ great grandparent). *NGG* = *N* ^*th*^ Great Grandparent. *ybp* = years before present, assuming 30 years per generation.

Although genetic material is passed down through a pedigree, Figure 2 shows that one can conceptualize IBD transmission as occurring upward. IBD transmission can occur either upward or downward. One can also conceptualize IBD transmission as occurring both upward and downward, for example between cousins.

### 3.2. The sampling approach

To sample conditional on the event that *i* and *j* share at least one IBD segment of at least τ cM with one another, we make use of Gibbs sampling in which we sequentially sample each meiosis in turn, conditional on all the other meioses. We first consider the case of transmission for a single chromosome and then generalize to the whole genome.

For meiosis *m*, let 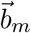 be the sorted list of recombination breakpoints among the two parental haplotypes undergoing recombination for a particular chromosome and let *h*_*m*_ ∈ {0, 1} be the parental haplotype on which copying starts. In a diagram of the pedigree, we adopt the convention that *h*_*m*_ = 0 corresponds to the haplotype of the “left” grandparent and *h*_*m*_ = 1 corresponds to the haplotype of the “right” grandparent. For example, in the meiosis between child *c* and parent *p* in Figure 1, haplotype 0 in *p* is the one inherited from *x*, whereas haplotype 1 in *p* is the haplotype inherited from *y*. In general, the lexicographically first parent label corresponds to the “left” parent.

Our approach is to use Gibbs sampling to sample the list 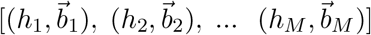 of all *M* = *u* + *d* + 2(*a* − 1) meioses between a pair of individuals by sampling each tuple 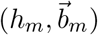 sequentially, one at a time, conditional on the breakpoints sampled at all other meioses.

For any given meiosis m, we use rejection sampling to sample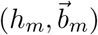. We sample h_*m*_ from a Bernoulli random variable. We then sample the number 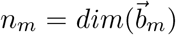of breakpoints from a Poisson random variable with mean *L*/100 and we sample the positions of each of the n_*m*_ breakpoints uniformly on the interval [0, *L*], where *L* is the length of the chromosome in cM. This approach is analogous to that of^24^ for the related problem of sampling breakpoints conditional on observed genotype data; however, instead of sampling the descent graph locus-by-locus along the genome, we sample all breakpoints at once and instead of conditioning on the exact observed genotypes (of which there are generally none) we condition on the event that specified pairs of individuals share IBD.

We first initialize the Gibbs sampler by sampling exactly one instance of 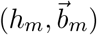 at each meiosis m without conditioning on observed IBD. We then proceed one meiosis at a time through the pedigree and we resample 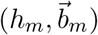 at each meiosis m conditional on the values 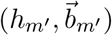 at all other meioses *m*^*′*^. We sample 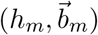 conditionally on the other meioses by proposing samples from the unconditional sampling distribution and rejecting proposals until we obtain a sample in which segments transmitted from *i* overlap by at least τ cM with segments transmitted from *j*. Once we achieve this result, we accept the proposed tuple 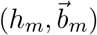 and move on to sample 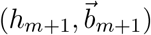.

#### 3.2.1. Acceptance rules when a segment can be transmitted

To illustrate the acceptance rule, a single meiosis in shown in Figure 3. In this meiosis, a child haplotype *c* inherits from the two haplotypes in parent *p* corresponding to parents *x* and *y* of *p*. We are interested in sampling the starting haplotype *h* and the breakpoints 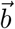 conditional on the event that some IBD is transmitted between relatives *i* and *j*.

**Figure 3.**
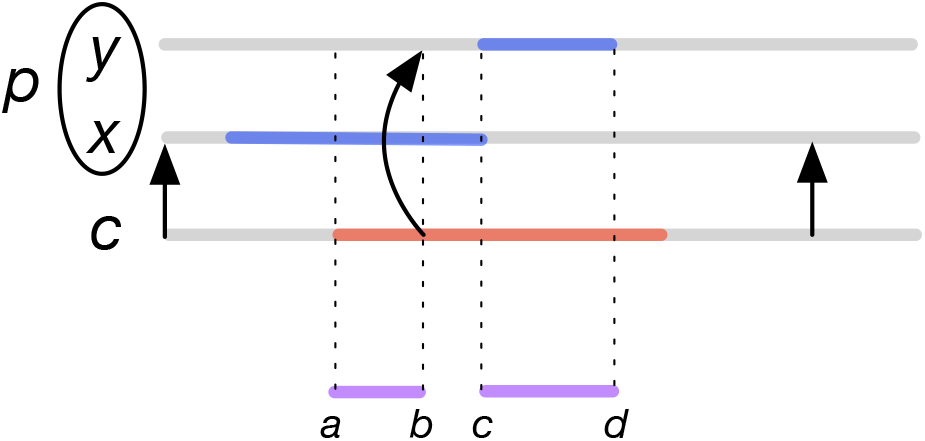
IBD propagation in a meiosis. An example of the meiosis between child node c and parent node *p* in Figure 1 is shown. The haplotype that *c* inherits from *p* is shown, along with the “left” (*x*) and “right” (*y*) haplotypes in individual *p*. The red region is the IBD tract that *c* shares with *i* and the blue regions are the tracts that *p* shares with *j*. Arrows indicate the haplotype that *c* copies from. From left to right, *c* begins copying on haplotype x of *p* (left-most arrow). It then switches to haplotype *y* (second arrow) and then back to *x* (third arrow). In the region between the vertical dashed lines *a* and *b*, individuals *i* and *j* both share IBD with haplotype x of *p*. In the region between vertical lines *b* and *c*, individual *i* shares with haplotype *y* and individual *j* shares with haplotype *x*. Between *c* and *d*, both *i* and *j* share IBD with haplotype *y*.

In Figure 3, the red region corresponds to genetic material in individual *c* that is shared with relative *i*. The blue region corresponds to genetic material in parent *p* that is shared with individual *j*. Note that *i* and *j* can share IBD with both haplotypes of *p*. A meiosis 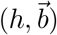 successfully transmits IBD between *i* and *j* if the red segments in *c* overlap with the blue segments in *p* by at least τ cM on a haplotype that is transmitted from *p* to *c*. These transmitted haplotypes are shown in purple in Figure 3. When the minimum length of a purple segments in Figure 3B is greater than τ, we accept the proposed tuple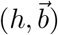.

A version of this sampling approach can also be used to sample from a genetic map. We discuss the genetic map approach in Section 3.4 and we discuss how to use the genetic map approach to sample from the full genome in Section 3.5.

#### 3.2.2. Acceptance rules when multiple meioses must be sampled together

In some cases, IBD between two individuals can be propagated along two different paths in the pedigree. A common example of this case is when two individuals share IBD through a pair of common ancestors. This scenario shown in Figure 1, in which *i* and *j* can share IBD either along the path through common ancestor *x* or common ancestor *y*, or through both.

In Figure 1, we cannot apply an acceptance criterion that requires IBD to be transmitted between *i* and *j* separately to each of the meioses (*p, x*) and (*p, y*). If we did so, we would require IBD to be transmitted along *both* paths, which is more restrictive than the criterion that IBD is transmitted at all. In such a case we must jointly sample both meioses (*p, x*) and (*p, y*) and accept a proposed sample as a success if either (*p, x*) or (*p, y*) is a success.

Moreover, it can dramatically increase the rate of acceptance if we jointly sample all meioses (*c, p*), (*p, x*), (*p, y*), (*z, x*), (*z, y*), and (*j, z*), which ensures that IBD from *i* and *j* is passed to the proper parental haplotypes in *x* and *y*. This kind of block Gibbs sampling is analogous to that implemented by Tong and Thompson ^24^ for a similar problem. If we did not include *c* and *j* in the sampling block, then we could accept meioses in which all segments from *i* are shared on the haplotype of *p* that comes from *y* and all segments from *j* are shared on the haplotype of *z* that comes from *x*. In such a case, we would not be able to sample meioses (*p, x*), (*p, y*), (*z, x*), and (*z, y*) in which segments from *i* are shared with *j* through *p* and *z*.

In general, sampling is most efficient if we jointly sample all meiosis pairs that form a block in which constraints must be simultaneously satisfied. Such blocks can be assembled by the following rules:

1. If meiosis (*c, p*) is in the block then all meioses involving full siblings of *c* must be in the block.
2. If meiosis (*c, p*) is in the block then all meioses between *p* and children of *p* must be in the block.

More general forms of block sampling can be used to handle more complicated loops. In such blocks, we jointly sample all meioses along all divergent paths in the loop. In principle, this would allow us to sample from inbred pedigrees, although we do not investigate inbreeding here. The longer the path, the more unlikely it is that IBD will be transmitted through it, making the simultaneous sampling of multiple paths increasingly less practical as the paths become longer.

#### 3.2.3. Acceptance rules when no segment can be transmitted

Note that in sampling meiosis (*c, p*) between child *c* and parent *p* in Figure 1, the other meioses in the pedigree may have been sampled in such a way that no segments from *i* have been propagated between *i* and *c* and/or no segments from *j* have been propagated between *p* and *j*. In such a case, we must establish rules for accepting or rejecting a proposed tuple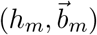).

Rules for accepting proposed samples when IBD cannot be transmitted are helpful for propagating IBD through the pedigree before we have reached a valid state in which IBD is shared among all pairs who must share IBD. These rules help us to arrive more quickly at a valid state.

When it is not possible to transmit any segment of at least τ cM we can still accept a tuple 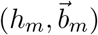 under the following conditions:

1. No segments are available either above or below to transmit; so transmission is impossible.
2. At least one IBD segment longer than τ cM is available from below to transmit and at least one IBD segment longer than τ cM is transmitted up.
3. At least one IBD segment longer than τ cM is available from above to transmit and at least one IBD segment longer than τ cM is transmitted down.
4. At least one IBD segment longer than τ cM is available both above and below to transmit and at least one IBD segment longer than τ cM is transmitted both down and up.

This rule for accepting proposed samples when it is not possible to fully transmit a segment longer than τ cM has the purpose of speeding up the transmission of IBD through the pedigree until all required IBD constraints have been transmitted. Once IBD has been transmitted among all IBD-sharing pairs, we can begin to accept proposed samples 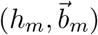 under the correct condition that at least one segment longer than τ cM is propagated.

In general, when sampling sets of meioses together, rather than one at a time, we apply the above criteria to the “receiving’ and “transmitting” nodes in the group, rather than the child and parent nodes. For instance, if our group consists of nodes *p, x, y* and *z* in Figure 1, then node *z* is the node in the group that “transmits” segments from *j* and node *p* is the node that “receives” these segments and passes them on to *i*. Thus, we require the following to be true in order to accept a sample jointly from (*p, x*), (*p, y*), (*z, x*) and (*z, y*):

1. If nodes *i* and *p* share at least one IBD segment longer than τ cM, then at least one IBD segment longer than τ cM must be transmitted to *z* (for propagation efficiency).
2. If nodes *j* and *z* share at least one IBD segment longer than τ cM, then at least one IBD segment longer than τ cM must be transmitted to *p* (for propagation efficiency).
3. If nodes *i* and *p* share at least one IBD segment longer than τ cM and if *j* and *z* share at least one IBD segment longer than τ cM and if at least one IBD segment longer than τ cM overlaps between these segments on a meiosis path (black or orange in 1), then at least one IBD segment of length τ cM of this overlap must be transmitted. Note that this criterion requires the overlap to occur on a feasible meiosis path (black or orange in Figure 1). If a segment in *z* physically overlaps a segment in *p*, but on opposite parental haplotypes, then the transmission is impossible and we must accept the proposed sample.

### 3.3. Data structures

In order to establish constraints on each meiosis, we set up and update several data structures. The first data structure, `ibd_dict`, records the segments propagated at each sampling step between each genotyped node in the pedigree and each haplotype of each other node in the pedigree. This dictionary is updated at each sampling step. Another data structure, `constraint_dict` records the pairs of genotyped IDs that must transmit IBD through each haplotype of each other node in the pedigree, along with the direction of the transmission; for example, the dict specifies that IBD is passed from node *i* to node *j* “up” through haplotype 0 of node *n*. This dictionary is established before sampling begins and does not change.

Another two dictionaries, `start_hap_dict` and `bpt_pos_list_dict`, store the starting haplotypes and breakpoint positions in each meiosis. These dictionaries are updated on each sampling step. Finally, the pedigree structure itself is stored in a dictionary, `up_dict` of the form {*c* : {*p*_1_, *p*_2_}, …} mapping each child node c to zero, one or two parent nodes. It is also convenient to store the reverse (`down_dict`) of `up_dict` mapping parents to their child nodes.

### 3.4. Sampling with a genetic map

Let 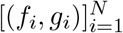 denote a genetic map from physical positions (denoted *f*_*i*_ since we used *p* for parents) to genetic positions *g*_*i*_. The values *f*_*i*_ are strictly increasing in *i* and the map from *f* to *g* is monotonically non-decreasing. To sample from such a map, we note that the genetic distance from *f*_*i*_ to *f*_*i*+1_ is *g*_*i*+1_ −*g*_*i*_. Therefore, we sample the number of breakpoints in the region (*f*_*i*_, *f*_*i*+1_) from a Poisson distribution with mean (*g*_*i*+1_ − *g*_*i*_)/100. The positions of these breakpoints in the region are then uniformly distributed over the interval (*f*_*i*_, *f*_*i*+1_). We sequentially sample in each region from left to right along the chromosome. The starting haplotype is sampled according to a Bernoulli distribution as usual.

### 3.5. Sampling a full genome with any transmitted IBD

Sampling a full genome is more complicated than sampling each chromosome independently because some IBD is guaranteed to be transmitted using our method. Thus, applying it to each chromosome separately would yield IBD on each chromosome. A simple work-around to this problem is to sample from the whole genome all at once by modeling it as a single, long chromosome amounting to all chromosomes concatenated end-to-end with gaps between them. Over this long chromosome, we apply a mask to ignore sampled breakpoints within the regions between chromosomes. We also set up our genetic map so that the gaps between chromosomes correspond to long genetic distances, thereby allowing essentially free recombination in these regions (Figure 4).

**Figure 4.**
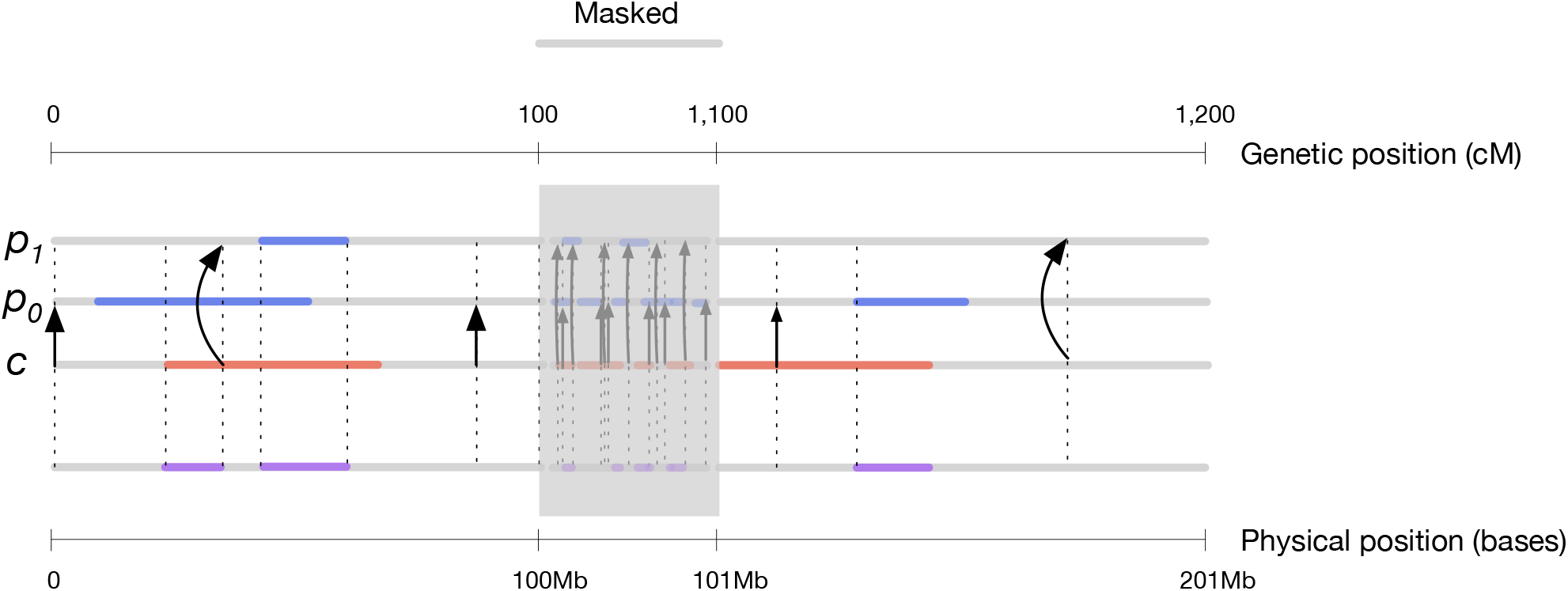
Masking. In the figure, two chromosomes, each of physical length 100MB and genetic length 100CM are sampled by simulating one long chromosome comprised of the two chromosomes concatenated together with a gap between them (region shaded by the grey rectangle). We ignore all segments and transmissions in the gap and only accept transmissions that occur outside of it. Regardless of the physical length of the gap, the genetic length of the gap can be set to a large number to approximate free recombination between the two chromosomes. In the figure, the gap is 1Mb long in physical coordinates, but 1,000 cM long in genetic coordinates. The approach can be extended to arbitrarily many chromosomes with the caveat that sampling in the masked regions requires compute time.

Another more efficient approach is to sample all chromosomes at the same time in a meiosis (i.e., without concatenating and masking) and accept a set of proposed breakpoints if at least one IBD segment of at least τ cM is transmitted in the full genome. This approach has the advantage that we do not need to sample segments within masked regions. Ultimately, either approach is efficient since we ignore masked segments. We present masking here because it is useful for ignoring chromosomal regions, such as those with high levels of background IBD or “pile-up.”

### 3.6. Comparison with rejection sampling

To check that the conditional sampler is producing the correct distribution, we compared it with simple rejection sampling for scenarios in which rejection sampling was computationally tractable. Figure 5 shows a comparison between the Gibbs conditional sampler (orange) and the simple rejection sampler (blue) for several different relationships. The simulations are based on one chromosome of length 100 cM. Each panel shows a histogram based on 1,000 samples. For each sample, the Gibbs sampler made 10 passes, sampling each group of meioses ten times. The event on which we conditioned was τ > 0 cM: i.e., that any IBD was transmitted between the two relatives.

**Figure 5.**
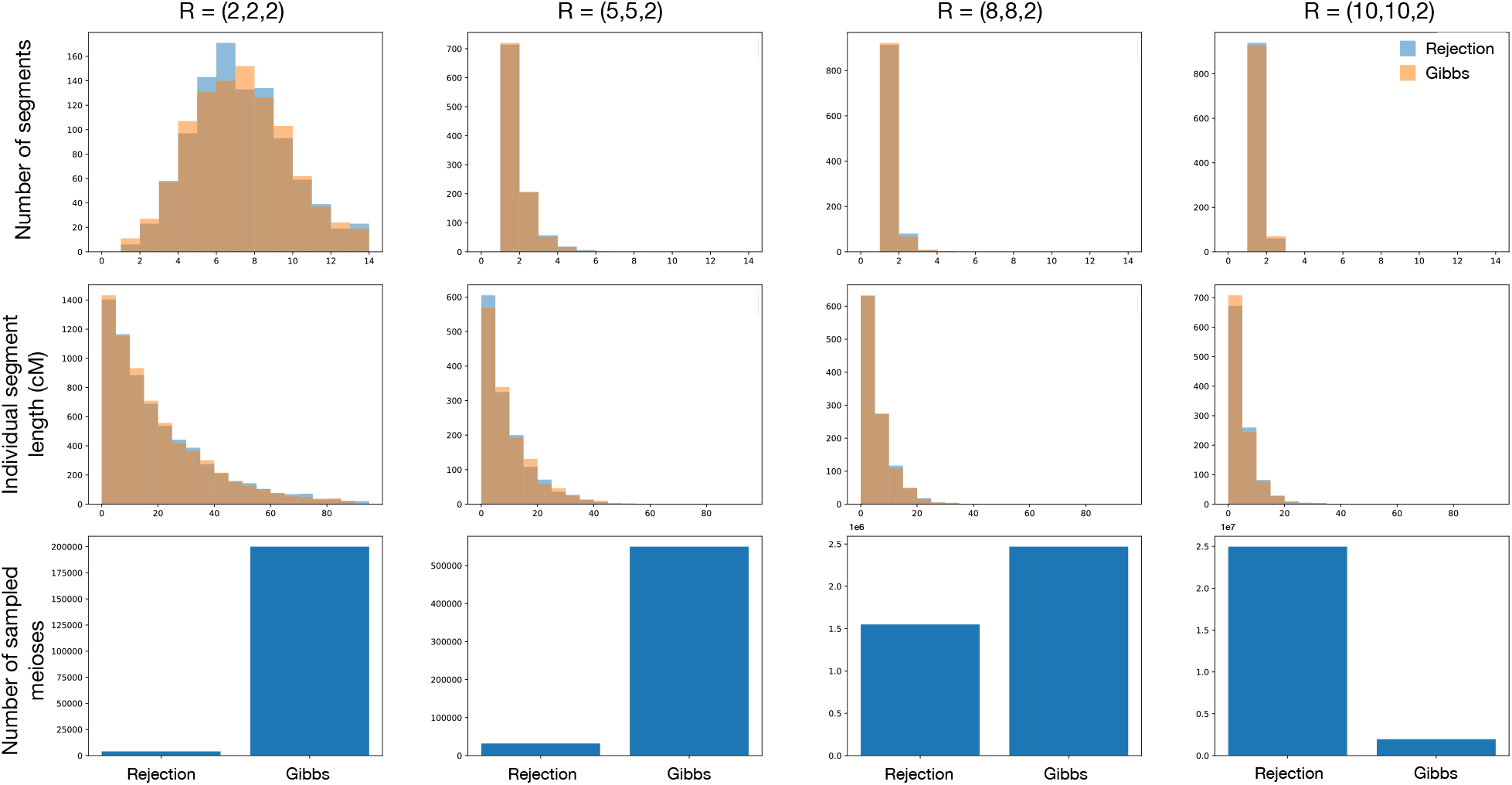
Comparison of the simple rejection sampler with the Gibbs sampler for different relationships *R* = (*u, d, a*). 1,000 replicates of each relationship were sampled using each method. Both samplers conditioned on the event that τ > 0 cM of IBD was transmitted between the two relatives.

Figure 5 shows that the conditional Gibbs sampler produces the same segment count and length distributions as the simple rejection sampler, but that the relative number of Gibbs replicates becomes increasingly small relative to the number of simple rejection replicates as the degree of the relationship becomes larger. From Figure 6, it can be seen that the overall timing for the Gibbs sampler stays manageable even beyond 30 generations in the past, at which point the probability of observing IBD between two genealogical relatives becomes low.

**Figure 6.**
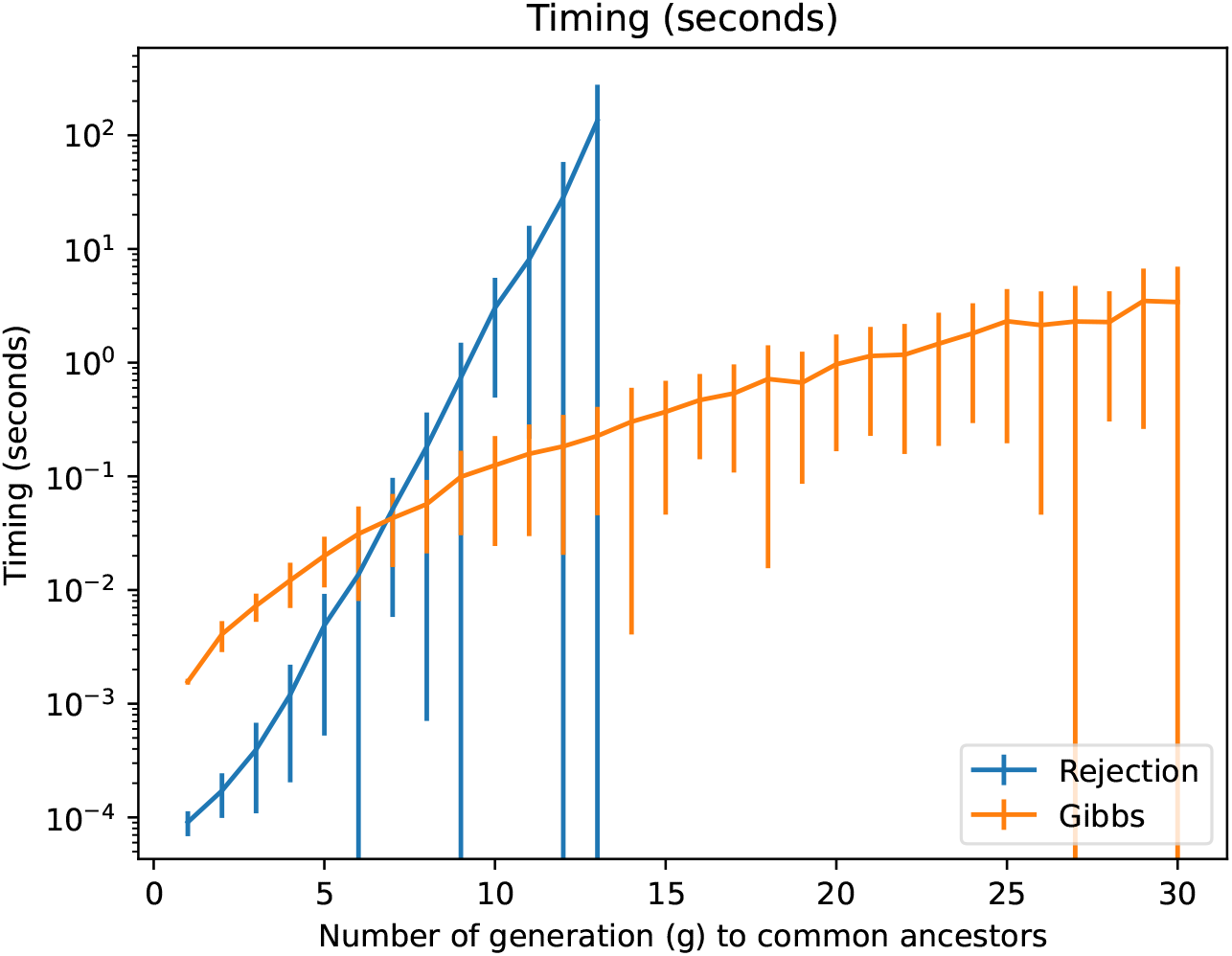
Timing of the simple rejection sampler compared with timing of the Gibbs sampler. Simulations were performed on a pair of *n*^*th*^ full cousins whose common ancestors lived *g* generations in the past. Each datapoint is the mean time over 100 replicates and error bars show one standard deviation above and below the mean. Simulations were performed only for *g* ≤ 13 for the simple rejection sampler, after which point rejection sampling became difficult.

Although the number of required replicates is higher for the Gibbs sampler when the degree of the relationship is small, it quickly becomes small relative to the number of replicates required for full rejection sampling. The relative computation time of the two samplers is shown in Figure 6.

#### 3.6.1. Sampling on a pedigree

The Gibbs conditional sampling approach becomes especially useful when there is more than one pair of relatives who must share IBD with one another. In this case, the probability that all constrained pairs share IBD at once can become exceedingly small, making simple rejection sampling impractical even for small pedigrees.

Figure 7 shows a comparison of the simple rejection sampler and the Gibbs conditional sampler for the pedigree structure in Figure 7A. In contrast with the analysis shown in Figure 5, we now condition on the much more restrictive event that three different pairs of genotyped individuals share IBD with one another. Specifically, we condition on the event that *i* shares IBD with *j, j* with *k*, and *i* with *k*.

**Figure 7.**
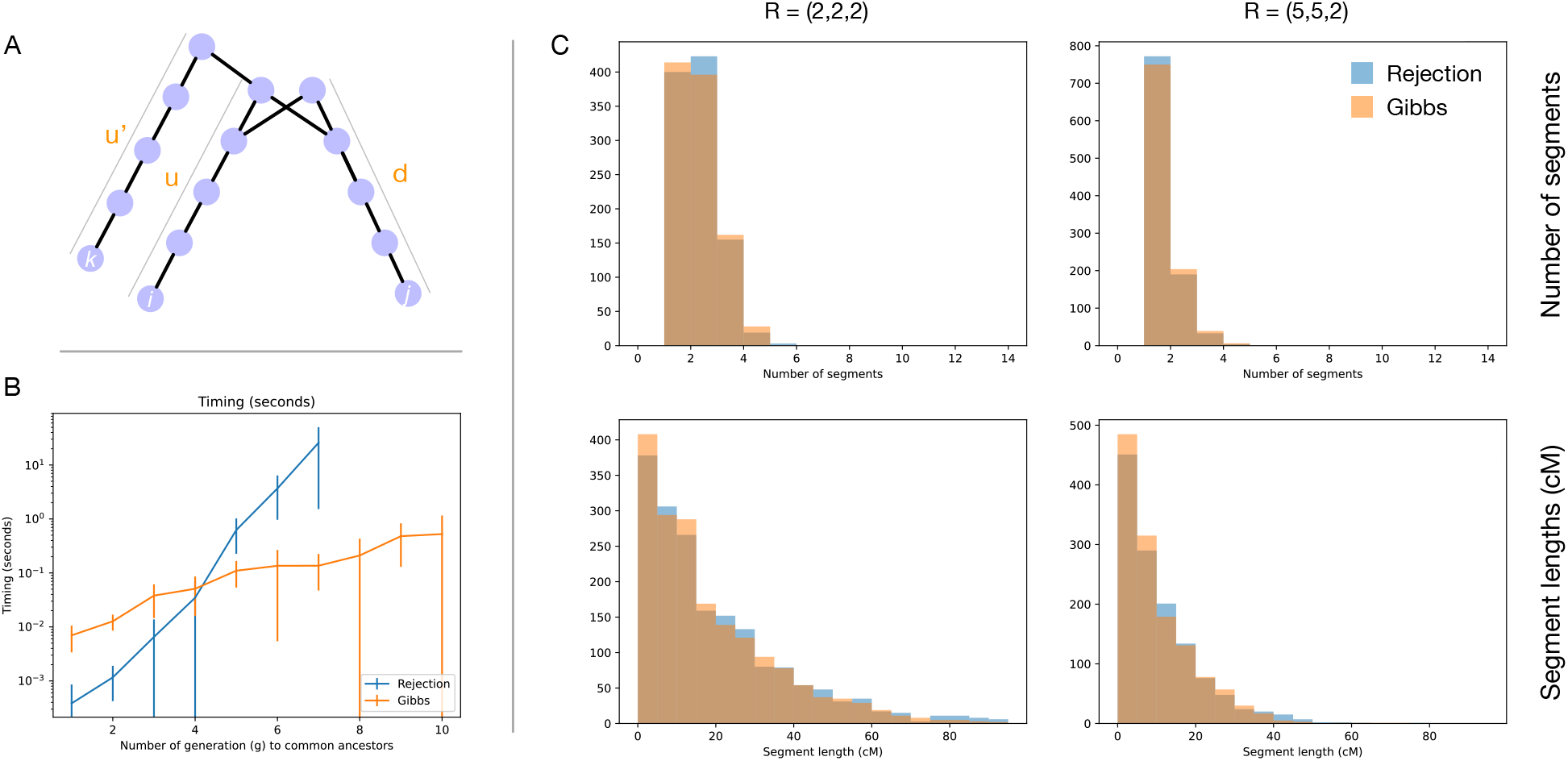
Comparison of rejection sampler and Gibbs sampler for a pedigree with multiple constrained pairs. (A) Pedigree structure. Individuals *i, j* and *k* are related such that *i* and *j* share two common ancestors and *k* shares a single common ancestor with each of *i* and *j*. (B) Timing for the rejection and Gibbs samplers for different values of *u, u*^*′*^, and *d*. In each pedigree, we constrained *u*^*′*^ = 2 and *u* = *d* = *g* and we varied *g* from 1 to 10. (C) Comparison of the rejection and Gibbs sampling distributions. Two statistics, the number of segments and segment lengths, are shown between *i* and *j* for relationship types *R* = (2, 2, 2) (first cousins) and *R* = (5, 5, 2) (fourth cousins).

Figure 7C shows a comparison of the distributions of segment counts and lengths for individuals *i* and *j*. This is easier to visualize than the full joint distribution of segments shared between all pairs of genotyped nodes *i, j* and *k*. By comparing the segment count distribution for the relationship *R* = (2, 2, 2) in Figure 7C with the count distribution for this relationship in Figure 5, it can be seen that these are not the same distributions due to the additional constraint that i and j must share IBD with k as well as with one another.

By comparing Figure 7B with Figure 6, it can be seen that including the additional requirement that IBD is shared with a third individual increases the computational time of both the simple rejection sampler and the Gibbs conditional sampler, but that it affects the simple rejection sampler much more than the conditional sampler. In particular, the Gibbs sampler becomes computationally more efficient than the simple rejection sampler when the common ancestors are around *g* = 4 generations in the past, compared with *g* = 7 generations in the past for the two-relative case shown in Figure 6. In the more highly constrained case of a three-person pedigree, the simple rejection sampler becomes unwieldy by *g* = 7 generations in the past, compared to *g* = 13 generations in the two-person case. In contrast to the rejection sampler, the Gibbs sampler can be used for quite distant relationships.

Although one can run rejection simulations in parallel, the expected amount of shared IBD decreases exponentially with each degree of separation between two IBD-sharing individuals or with the addition of IBD-sharing pairs. As a result, one cannot simply run jobs in parallel to overcome the computational slowness of rejection sampling. For instance, the probability that two individuals share any IBD when they are separated through a single common ancestor who lived 15 generations in the past is approximately 5 × 10^*−*7^ and when the ancestor existed 20 generations in the past this probability^10^ is approximately 6 × 10^*−*10^.

Adding in additional constrained pairs quickly makes the problem completely intractable. For instance, the lineage extending upward from 1 to 16 via nodes −2 and −5 in Figure 9 is nearly independent of the lineage extending upward from nodes 1 to 10 through nodes −2 and −4. If we wish to jointly sample IBD sharing among nodes 1 and 16 and 1 and 10, we would need to satisfy these two nearly independent constraints, each of which has a probability^10^ of approximately 10^*−*3^. From this, we see that the sampling problem quickly becomes computationally challenging after the addition of a few IBD-sharing pairs.

## 4. Applications

### 4.1. Simulating IBD-sharing distributions for relationship inference

The conditional sampler described in Section 3 is essential for simulating IBD between distant relatives who are known to share some amount of IBD. An immediate application of this sampler is simulating IBD sharing distributions that can be used in relationship estimators. As we have noted, the unconditional distributions that are currently used for this purpose are inappropriate because they do not account for the fact that putative relatives are generally ascertained on the basis of IBD sharing with one another.

For relationships of increasingly distant degree, Figure 8A shows the mean total IBD between two relatives separated by various degrees. The mean IBD from the unconditional distribution is shown in blue. The IBD from the conditional distribution (requiring τ > 0) is shown in orange. From Figure 8A, it can be seen that the two means begin to diverge when relationships are approximately 8 degrees (approximately third cousins). For relationships ten degrees and greater (approximately fourth cousins) the distributions have largely diverged. The unconditional mean total IBD goes to zero quickly and cannot be shown on the log-scale plot in Figure 8A for degrees above approximately 20.

**Figure 8.**
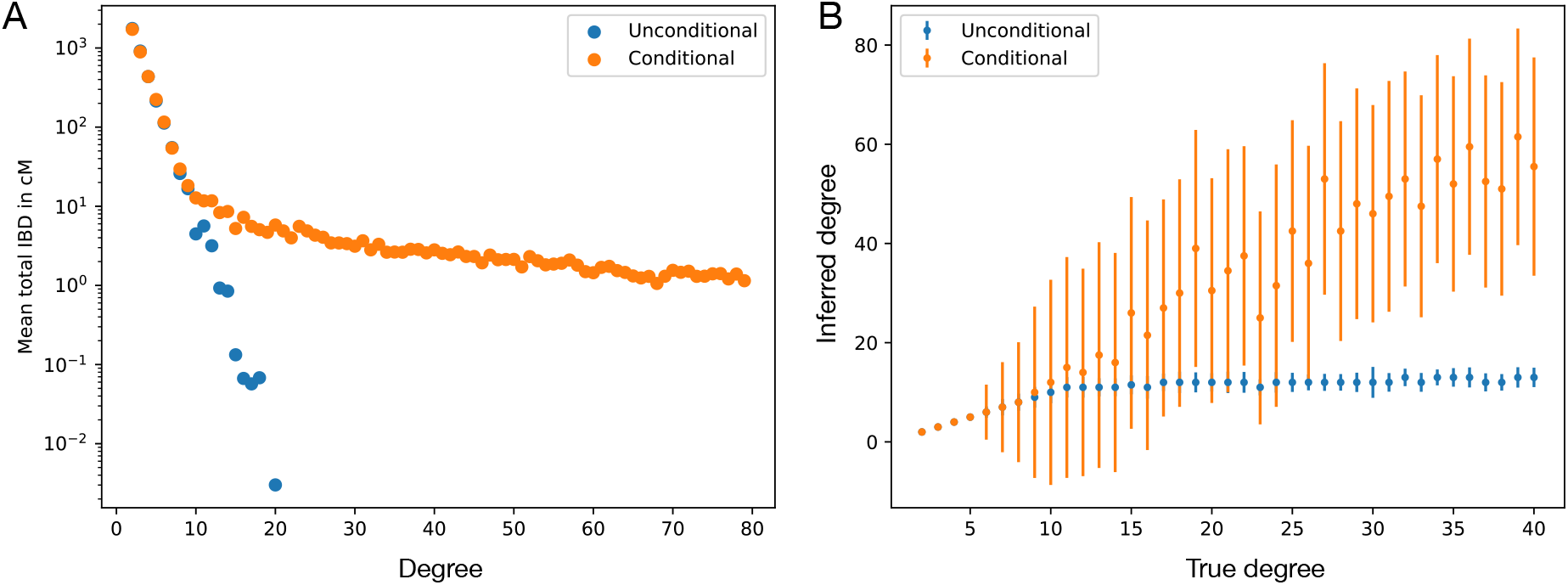
Comparison of the unconditional and conditional sampling distributions. (A) Mean total IBD as a function of degree between two individuals connected through a single common ancestor. (B) A simple method of moments estimator of degree applied to the total amount of IBD between relatives of different degrees.

**Figure 9.**
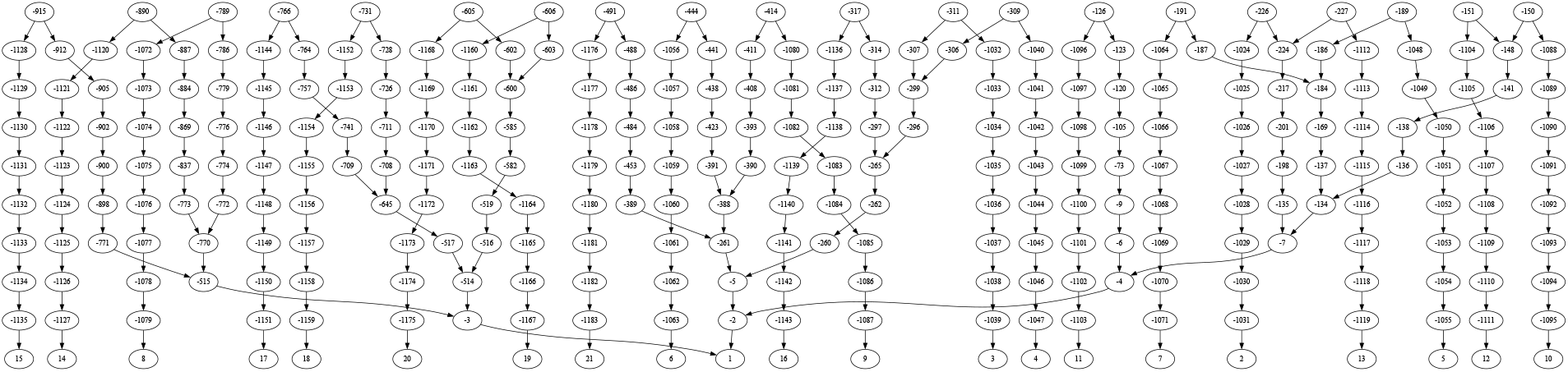
An example of a pedigree containing an individual and twenty of their distant relatives that can be simulated using the conditional sampler. Each individual is denoted by a disc and each line represents a parent-child connection between two individuals. Genotyped nodes are indicated with positive numbers and ungenotyped nodees are indicated with negative numbers. In this topology, a focal individual (1) is connected to their distant relatives, each through a single common ancestor who lived ten generations in the past.

The discrepancy between the conditional and unconditional distributions has major implications for relationship inference. Figure 8B shows degree estimates from a very simple method of moments estimator. The estimator takes the total IBD observed for a pair of individuals and finds the degree for which the expected total length of IBD most closely matches the observed total IBD. This kind of estimator is similar to those used by several direct-to-consumer genetic testing companies and bias engendered by employing the unconditional distribution is found in all currently-used relationship estimators.

Figure 8B shows that the estimator based on the unconditional distribution is heavily biased and reaches an asymptote near ten degrees. This explains the fact that the most distant relationships inferred by existing estimators are around ten degrees. In contrast, the unconditional estimator makes it possible to infer much greater degrees of relationship, although the moment estimator is still biased. The implication of Figure 8 is that one must use conditional sampling to obtain empirical distributions that are used for relationship inference in the usual case in which putative relatives are ascertained on the basis of IBD sharing.

### 4.2. Simulating the pedigrees of today

In datasets that contain many genotyped or sequenced individuals like the UK Biobank ^7^ and direct-to-consumer genetic testing databases ^2,9^, genealogical relationships can be detected that extend tens of generations into the past.

There is a strong need to be able to simulate data for these kinds of pedigrees in order to support the development of new algorithms. The conditional sampler presented in this paper makes it possible to simulate IBD between an individual and a set of distant relatives.

Figure 9 shows one such pedigree, including an individual and twenty of their distant relatives, each separated from the focal individual by 20 degrees of relatedness. This pedigree was simulated in under a minute using the conditional sampler. A single lineage of this pedigree would require an average of 10^3^ replicates to achieve an acceptance from the rejection sampler and all 20 lineages together would require approximately 10^60^ replicates, rendering the simulation effectively impossible using existing simulation methods.

## 5. Discussion

Methods like the one presented here are necessary in order to simulate the kinds of pedigrees that appear in large genotyping datasets. As these databases increase in size, simulation methods will need to handle increasingly large and dense pedigrees.

Several aspects of the existing simulator can be improved in order to decrease the runtime. In particular, we have been somewhat cavalier about the rejection sampler that is still employed to sample each meiosis in the Gibbs sampler. This rejection sampler does not take into account the regions in which sampled breakpoints must or must not occur and therefore it can be fairly wasteful. For example, in the pedigree in Figure 9, we might find that the tract transmitted from node 6 (the node to the immediate left of node 1) is in close proximity to the tract transmitted from node 16 (the node to the immediate right of node 1). Since these tracts must lie on opposite haplotypes in ancestor −5 of node 1 (though potentially near each other on the same chromosome), we must achieve a recombination event between them. If the distance δ between these tracts is small, the probability of obtaining a recombination event in this region will be small. In order to speed up sampling, we can constrain the locations in which breakpoints must or must not occur.

The existing sampler makes it possible to simulate from genetic maps including sex-specific genetic maps. However, it is important to account for additional phenomena like cross-over interference ^3^. Future versions of this sampling method will be updated to include cross-over interference.

Using IBD sharing statistics simulated with this approach, it is possible to replace the biased distributions that are still used for relationship inference with distributions that are appropriate for the inference problem. Doing so has a considerable effect on the distant relationships inferred in large genotyping databases, moving them from five or six generations in the past to fifteen or twenty generations in the past or more. The variance in these estimates also increases substantially, reflecting true underlying uncertainty.

## 6. Acknowledgements

I would like to thank the employees and research participants of 23andMe who made this research possible. I am especially grateful to Amy L. Williams, William A. Freyman and David A. Hinds for their insightful comments and thoughtful reviews of this manuscript and Peter R. Wilton for insightful points raised in discussions. Funding for this work was provided by NIH grant R35 GM133805 and by 23andMe, Inc. Members of the 23andMe Research Team are Stella Aslibekyan, Adam Auton, Elizabeth Babalola, Robert K. Bell, Jessica Bielenberg, Ninad S. Chaudhary, Zayn Cochinwala, Sayantan Das, Emily DelloRusso, Payam Dibaeinia, Sarah L. Elson, Nicholas Eriksson, Chris Eijsbouts, Teresa Filshtein, Pierre Fontanillas, Davide Foletti, Will Freyman, Zach Fuller, Julie M. Granka, Chris German, Éadaoin Harney, Alejandro Hernandez, Barry Hicks, David A. Hinds, M. Reza Jabalameli, Ethan M. Jewett, Yunxuan Jiang, Sotiris Karagounis, Lucy Kaufmann, Matt Kmiecik, Katelyn Kukar, Alan Kwong, Keng-Han Lin, Yanyu Liang, Bianca A. Llamas, Aly Khan, Steven J. Micheletti, Matthew H. McIntyre, Meghan E. Moreno, Priyanka Nandakumar, Dominique T. Nguyen, Jared O’Connell, Steve Pitts, G. David Poznik, Alexandra Reynoso, Shubham Saini, Morgan Schumacher, Leah Selcer, Anjali J. Shastri, Jingchunzi Shi, Suyash Shringarpure, Keaton Stagaman, Teague Sterling, Qiaojuan Jane Su, Joyce Y. Tung, Susana A. Tat, Vinh Tran, Xin Wang, Wei Wang, Catherine H. Weldon, and Peter Wilton.

## References

[1] G.R. Abecasis, S.S. Cherny, W.O. Cookson, and L.R. Cardon. Merlin-rapid analysis of dense genetic maps using sparse gene flow trees. Nat. Genet., 30:97–101, 2002.

[2] C.A. Ball, M.J. Barber, J. Byrnes, P. Carbonetto, K.G. Chahine, R.E. Curtis, J.M. Granka, E. Han, E.L. Hong, A.R. Kermany, N.M. Myres, K. Noto, J. Qi, K. Rand, Y. Wang, and L. Willmore. Rapid forward-in-time simulation at the chromosome and genome level. https://www.ancestry.com/dna/resource/whitePaper/AncestryDNA-Matching-White-Paper.pdf, 2016.

[3] M. Caballero, D.N. Seidman, Y. Qiao, J. Sannerud, T.D. Dyer, D.M. Lehman, J.E. Curran, R. Duggirala, J. Blangero, S. Carmi, and Williams A.L. Crossover interference and sex-specific genetic maps shape identical by descent sharing in close relatives. PLoS Genet., 15:e1007979, 2019.

[4] R.H. Chung and C.C. Shih. SeqSIMLA: a sequence and phenotype simulation tool for complex disease studies. BMC Bioinformatics, 14:199, 2013.

[5] A. Dimitromanolakis, J. Xu, A. Krol, and L. Briollais. sim1000G: a user-friendly genetic variant simulator in R for unrelated individuals and family-based designs. BMC Bioinformatics, 20:26, 2019.

[6] K. P. Donnelly. The probability that related individuals share some section of genome identical by descent. Theoretical Population Biology, 23(1):34–63, Feb 1983. doi: 10.1016/0040-5809(83)90004-7.

[7] P. Downey and T.C. Peakman. Design and implementation of a high-throughput biological sample processing facility using modern manufacturing principles. International Journal of Epidemiology, 37:i46–i50, 2008.

[8] K. Finke, M. Kourakos, G. Brown, H.T. Dang, S.J.S Tan, Y.B. Simons, S. Ramdas, A.A. Schäffer, R.L. Kember, M. Bućan, and S. Mathieson. Ancestral haplotype reconstruction in endogamous populations using identity-by-descent. PLoS computational biology, 17:e1008638, 2021.

[9] B.M. Henn, L. Hon, J.M. Macpherson, N. Eriksson, S. Saxonov, I. Pe’er, and J.L. Mountain. Cryptic distant relatives are common in both isolated and cosmopolitan genetic samples. PLOS One., 7:e34267, 2012.

[10] C.D. Huff, D.J. Witherspoon, T.S. Simonson, J. Xing, W.S. Watkins, Y. Zhang, T.M. Tuohy, D.W. Neklason, R.W. Burt, S.L. Guthery, S.R. Woodward, and L.B. Jorde. Maximum-likelihood estimation of recent shared ancestry (ersa). Genome Research, 21:768–774, 2011.

[11] E.M. Jewett, K.F. McManus, W.A. Freyman, and A. Auton. Bonsai: An efficient method for inferring large human pedigrees from genotype data. Am. J. Hum, Genet., 108:2052–2070, 2021.

[12] Steven T. Kalinowski and Philip W. Hedrick. An improved method for estimating inbreeding depression in pedigrees. Zoo Biology, 17(6):481–497, 1998. doi: 10.1002/(SICI)1098-2361(1998)17:6⟨481::AID-ZOO2⟩3.0.CO;2-G. URL https://onlinelibrary.wiley.com/doi/abs/10.1002/%28SICI%291098-2361%281998%2917%3A6%3C481%3A%3AAID-ZOO2%3E3.0.CO%3B2-G.

[13] J. Kelleher, K.R. Thornton, J. Ashander, and P.L. Ralph. Efficient pedigree recording for fast population genetics simulation. PLoS Comput. Biol., 14:e1006581, 2018.

[14] A. Ko and R. Nielsen. Composite likelihood method for inferring local pedigrees. PLOS Genet., 13:e1006963, 2017.

[15] B. Li, G. Wang, and S.M. Leal. SimRare: a program to generate and analyze sequence-based data for association studies of quantitative and qualitative traits. Bioinformatics, 28:2703–2704, 2012.

[16] B Li, G.T. Wang, and Leal S.M. Generation of sequence-based data for pedigree-segregating Mendelian or Complex traits. Bioinformatics, 15:3706–3708, 2015.

[17] Mark P. Miller, Susan M. Haig, Jonathan D. Ballou, and E. Ashley Steel. Estimating Inbreeding Rates in Natural Populations: Addressing the Problem of Incomplete Pedigrees. Journal of Heredity, 108(5):574–582, 05 2017. ISSN 0022-1503. doi: 10.1093/jhered/esx032. URL https://doi.org/10.1093/jhered/esx032.

[18] C. Nieuwoudt, A. Brooks-Wilson, and J. Graham. SimRVSequences: an R package to simulate genetic sequence data for pedigrees. Bioinformatics, 36:2295–2297, 2019.

[19] Ying Qiao, Ethan M. Jewett, Kimberly F. McManus, William A. Freyman, Joanne E. Curran, Sarah Williams-Blangero, John Blangero, The 23andMe Research Team, and Amy L. Williams. Reconstructing parent genomes using siblings and other relatives. bioRxiv, 2024. doi: 10.1101/2024.05.10.593578.

[20] M.D. Ramstetter, S.A. Shenoy, T.D. Dyer, D.M. Lehman, J.E. Curran, R. Duggirala, J. Blangero, J.G. Mezey, and A.L. Williams. Inferring identical-by-descent sharing of sample ancestors promotes high-resolution relative detection. Am. J. Hum. Genet., 103:30–44, 2018.

[21] Schäffer, A.A. and Lemire, M. and Ott, J. and Lathrop, G.M. and Weeks, D.E. Coordinated conditional simulation with slink and sup of many markers linked or associated to a trait in large pedigrees. Hum. Hered., 71:126–134, 2011.

[22] J. Staples, D. Qiao, M.H. Cho, E.K. Silverman, University of Washington Center for Mendelian Genomics, D.A. Nickerson, and J.E. Below. PRIMUS: Rapid reconstruction of pedigrees from genome-wide estimates of identity by descent. Am. J. Hum. Genet., 95:553–564, 2014.

[23] J. Staples, D.J. Witherspoon, L.B. Jorde, D.A. Nickerson, University of Washington Center for Mendelian Genomics, J.E. Below, and C.D. Huff. PADRE: Pedigree-aware distant-relationship estimation. Am. J. Hum. Genet., 0:10.1101/2020.02.25.965376, 2016.

[24] L. Tong and E. Thompson. Multilocus Lod scores in large pedigrees: combination of exact and approximate calculations. Hum. Hered., 65:142–153, 2007.

[25] R.E. Voorrips and C.A. Maliepaard. The simulation of meiosis in diploid and tetraploid organisms using various genetic models. BMC Bioinformatics., 13:248, 2012.

[26] A.L. Williams. 2024. URL https://hapi-dna.org/2020/11/how-often-do-two-relatives-share-dna-2/.

